# The Devil Is In The Details: Landscape Features Are Insufficient To Explain Patterns Of Non-Native Fishes Distribution In North Patagonian Streams

**DOI:** 10.1101/2022.04.24.489319

**Authors:** Mailén Elizabeth Lallement, Magalí Rechencq, Eduardo Enrique Zattara

## Abstract

Ecological communities are structured by combinations of factors known as habitat templates. These templates work as a filter allowing only species with particular traits or phenotypes to establish and persist excluding all others. Defining which habitat variables and spatial scales drive the assembly of freshwater communities is key to effective and efficient management of fluvial ecosystems. We took advantage of the relatively recent and well-studied history of salmonid introductions in Patagonia to evaluate if non-native species show different patterns of association with abiotic factors depending on the spatial scale of the environmental filter. We used a hierarchical approach to characterized environmental variables at the basin and reach scales to assess their influence on the presence, abundance and structure of the salmonid assemblages in breeding streams. We saw no evidence that presence/absence patterns of salmonid distribution were driven by landscape variables, except for those basins with physical environmental barriers to colonization. However, we did find evidence for relative abundances being influenced by climatic and geomorphological variables (e.g., precipitation and relief). Our results do not support a scenario in which any of the salmonid species modulates the distribution of the other species, suggesting that interference has played only a minor role in determining current fish distribution in fluvial systems of the region. Instead, current patterns of presence and abundance of salmonids are best explained as the product of environmental filters. Our findings contribute to our understanding of the ecology of individual species and provide insight into the mechanisms structuring fish assemblages in Southern Hemisphere’s lotic systems.

## Introduction

Ecological communities are structured by combinations of factors defined as habitat templates (Southwood, 1977). These habitat templates mold evolutionary forces and structure ecological strategies for each of the species within the community. In other words, both the presence of a species at a particular site and its life history strategy are the result of environmental filtering (i.e., the tolerance of the species to a particular subset of biotic and abiotic characteristics) and trade-offs met during habitat adaptation.

Fluvial ecosystems are characterized by a physical habitat strongly influenced by the inherent hierarchical structure and patchiness that determine the distribution of organisms, food availability, predation and competition (Frissell *et al*., 1986; Schlosser & Kallemeyn, 2000). Fine-scaled variability results from the interaction between large-scale landscape variables (e.g., basin area, slope profile of stream-associated valleys and other geomorphological traits) and smaller scale variables (e.g., local structure and condition) (Frissell *et al*., 1986; Schlosser & Kallemeyn, 2000). Because of that, defining which habitat variables and spatial scales have the most influence on freshwater communities is key to effective and efficient management of fluvial ecosystems (Matthews, 1998; Gido *et al*., 2006).

Environmental filtering is considered a major structuring mechanism of communities (Weiher & Keddy, 1995). It is dominated by three ecological factors: dispersal restriction, abiotic environment and biotic interactions (Belyea & Lancaster, 1999). The first two act on a regional scale and delimit the area of action of the third, which operates on a local scale (Booth & Swanton, 2002). While the utility of the environmental filtering concept has been disputed on the basis that it predicts patterns that cannot be distinguished from those produced by other mechanisms, such as competition (Kraft *et al*., 2015), there are good reasons to explore how patterns of trait or phylogenetic dispersion change in response to the environment (Cadotte & Tucker, 2017).Therefore, if the relative effects of these general classes of factors in streams could be disentangled, then we would gain a comprehensive view of how each factor drives community composition.

In Patagonia, particularly at the scale of large drainage basins, fish distribution has been strongly influenced by the Andean uplift and the Quaternary glacial cycles (Hubert & Renno, 2006). After the retreat of glaciers during the Pleistocene, the ability of Patagonian fish to colonize post-glacial water bodies determined their present distribution, clearly constrained by climate, especially by temperature (Ruzzante *et al*., 2006; Cussac *et al*., 2009). In recent times, native freshwater fish communities have also been altered on repeated occasions by the introduction of non-native species to generate sport fisheries, resulting in communities where up to six native species and four introduced salmonid species coexist within the watershed (Macchi *et al*., 1999). Many of these species have stable populations at several large lakes that have been intensively studied in the past years (Cussac *et al*., 2014). However, little is known about the original distribution of fish assemblages in streams or about the environmental filters that structure current riverine communities (Pascual *et al*., 2002, 2007; Aigo *et al*., 2008; Barriga *et al*., 2013; Lallement *et al*., 2020).

Of the four species introduced to continental Patagonia since 1904, rainbow trout (*Onchorhynchus mykiss*, Walbaum 1792), brown trout (*Salmo trutta*, Linnaeus 1758) and brook trout (*Salvelinus fontinalis*, Mitchill 1814) are currently the most widely distributed and abundant species. The rapid adaptation of these salmonid species to the new environment enabled them to establish self-sustaining populations, obviating the need for continuous import of eggs and ensuring of a steady supply of fish from local reproducers. The short and well-known history of introductions in the region shows these species have high dispersive capabilities and found practically no biological resistance to invasion (Pascual *et al*., 2007). Thus, it can be posited that their current distribution in Patagonia resulted mainly from environmental filtering at different scales. Although the influence of the landscape scale factors has been observed for the two most abundant species of salmonids in previous works (Aigo *et al*., 2008; Lallement *et al*., 2020), it is uncertain if these factors alone explain the richness of non-native species in the watershed.

The objective of this work was to examine watershed and reach-level patterns of introduced salmonid species in North Patagonian rivers and their association with watershed characteristics derived from remote sensing and topography data across an environmental gradient. We compared the relative variation in salmonid density across different kind of watersheds expecting that some environmental characteristics at a landscape and reach scale conditioned i) the presence of each salmonid species, ii) the abundance of each species, iii) the assemblage conformation and iv) the dominance of a species at a regional level. We characterized environmental variables at the basin and reach scale to assess their influence on the presence, abundance and structure of the salmonid assemblages in breeding streams of the Upper Limay river basin. Our findings contribute to our understanding of the ecology of individual species and provide insight into mechanisms structuring fish assemblages in Southern Hemisphere’s stream systems that could inform conservation and management activities.

## METHODS

### Study Area

The Upper Limay River basin is located between the provinces of Río Negro and Neuquén (∼40°63’ S, 71° 70’ W), Argentina, and drains an area of 6.980 km^2^, most of it within the boundaries of Nahuel Huapi National Park (Figure 1). Originating in the eastern slopes of the Andes mountain range, the basin presents a complex hydrological network, with many streams, rivers and lentic water bodies. Due to rain-shadowing effects by the Andes, the area experiences a steep longitudinal climatic gradient going from 3000 mm of yearly precipitation in the West to less than 600 mm to the East in about 60 km (Paruelo *et al*., 1998). This climatic gradient results in an eastward transition from a cold-temperate rainforest to shrubby dry steppes. The watershed has a highly connected, complex hydrologic network characterized by deep oligotrophic lakes of varying size, interconnected by streams, ponds and wetlands. The main hub of the network is Nahuel Huapi Lake, with an area of 529 km^2^ and a maximum depth of 464 m, which collects most waters from the basin, and drains through the Limay River towards the Atlantic Ocean (Figure 1).

**Figure 1.**
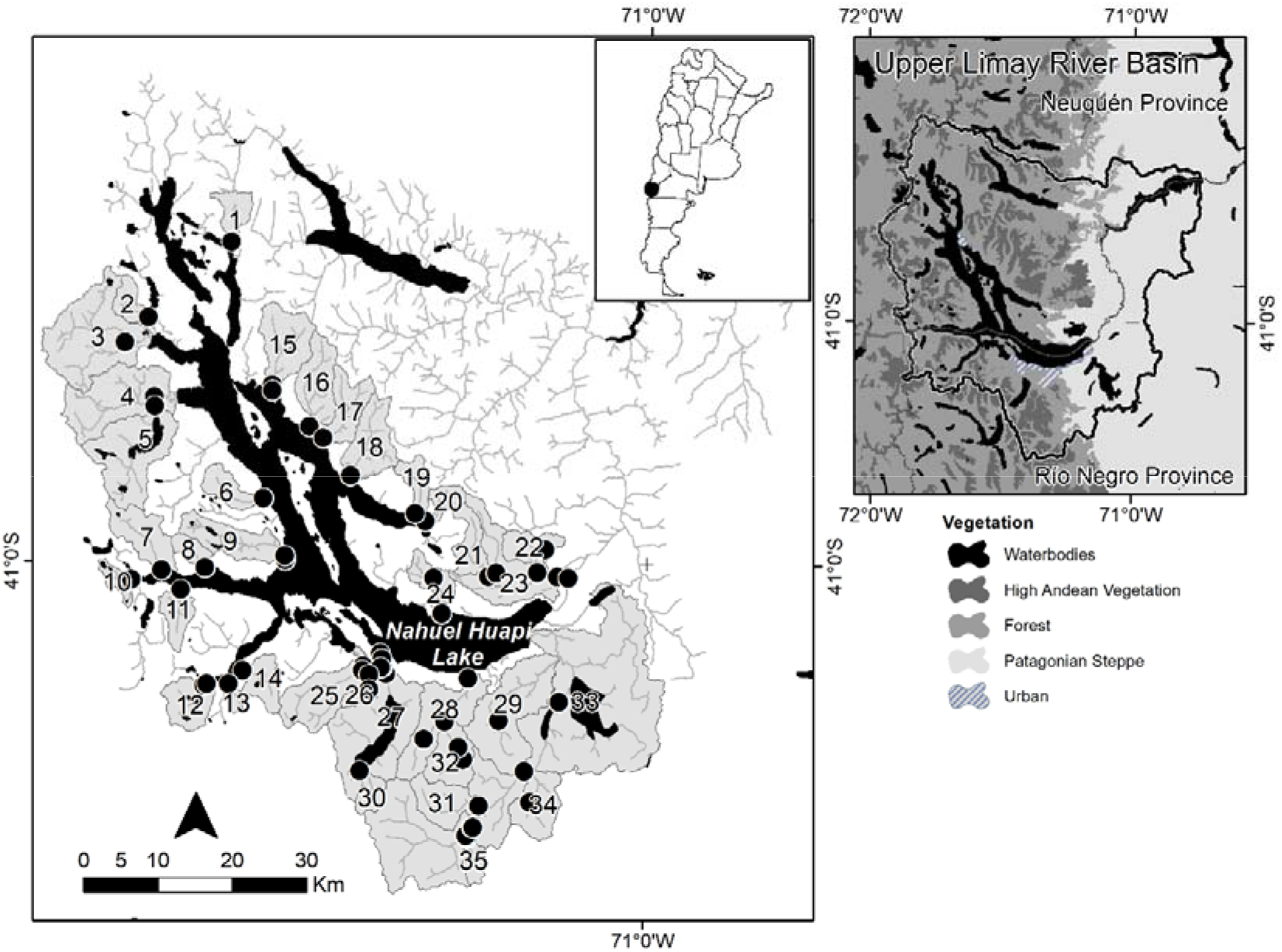
Sampling sites in each watershed in the Upper Limay river Basin: 1, Neuquenco; 2, Acantuco; 3, Pireco; 4, Machete; 5, Gallardo; 6, Coluco; 7, Bravo; 8, Blanco; 9, Millaqueo; 10, Blest; 11, Patiruco; 12, Frey; 13, Uhueco; 14, Lluvuco; 15, Bonito; 16, Estacada; 17, Ragintuco; 18, Huemul; 19, Pedregoso; 20, Quintriquenco; 21, Manzano-Jones; 22, Newbery; 23, Chacabuco; 24, Castilla; 25, Casa de Piedra; 26, Cascada; 27, Gutierrez; 28, Ñireco; 29, del Medio; 30, Torrontegui; 31, Tristeza; 32, Challhuaco; 33, Ñirihuau; 34, Las Minas; 35, de las Quebradas. Black dots are sampled reaches.

### Selection of watersheds

Thirty-five basins fully representing the environmental gradient of the watershed were selected for this study (Figure 1). For each basin, one or more sections of 2^nd^ order or higher were selected; selection criteria were size, registered physiographic changes, existence of natural or anthropic barriers and the accessibility to each section (Bain & Stevenson, 1999). Based on these criteria, some basins were represented by a single section, while others had two or more representative sections. On larger basins with enough altitudinal range (e.g., Machete, Gutiérrez, Ñireco, Ñirihauau and Chacabuco), additional sections on tributary streams were sampled.

### Watershed variables

A total of 32 watershed attributes were chosen following the available literature (Angermeier & Winston, 1999; Oakes *et al*., 2005; Smith & Kraft, 2005), and grouped into four distinct categories (Geomorphological, Climatological, Land Use and Vegetation) based on the general aspect of the environment measured by each variable (Tables AI,II,III-Appendix section). All categories were calculated using GIS tools (version 2.6) under the QGIS environment (QGIS Development Team, 2014), or using specific formulas following Bain and Stevenson (1999). Land Use and Vegetation data were obtained from existing digital map layers available from the National Geographic Institute, the National Institute of Agricultural Technology (INTA) and National Parks Administration’s Biodiversity Information System; these maps were complemented with satellite imagery from Google Earth. Watershed morphological variables were calculated based on a digital elevation model with a resolution of 30 m (Landsat 6TM+). Average annual precipitation (mm) and temperature were calculated based on a map interpolating averages of daily measures from 25 meteorological stations located within from 38°46’0S–41°30’0S and 70°03’0W–71°45’0W (Barros *et al*., 1983). The Normalized Differential Vegetation Index (NDVI) was calculated from Landsat 6TM+ satellite images during the summer season of 2014. Geoprocessing and zonal statistics were computed using Quantum Gis (QGIS) (version 2.6) and digital map data. All the variables are available in the Appendix section (Table A-I, II, III).

### Local Variables

Physicochemical variables were collected in 42 reaches. Variables included geomorphological, substrate type, hydrological, water quality and others (Table B-Appendix section). Location and number of sections sampled by stream were determined by the pattern shape of channels, covered area and accessibility, which determined that streams with the largest area have a greater number of sections sampled. Thus, sections were selected based on particular characteristics of each stream such as changes in the relief or riverine vegetation, tributary unions and presence of ponds or waterfalls.

### Fish Capture

Different stream segments that presented a succession of pool-riffle-pool habitats were sampled during the austral summer (from December to March) of 2013-2014. The minimum sampled area depended on the size of the selected reach. In each section, presence-absence data were collected by three-pass electrofishing with a Smith-Root mod 12B equipment along 50-m reach. Relative abundance, expressed as catch per unit effort (CPUEN), was standardized based on actual length of each sweep to number of fish caught per 100 m^2^. Electrofishing was conducted following an upstream zigzag trajectory and exploring all habitat types. The extremes of the trajectory were not blocked with nets. For basins sampled in more than one section, an average of the CPUEN of all sampled sections was used as basin-level fish abundance. The Administration of Nahuel Huapi National Park and San Carlos de Bariloche Town Council approved our protocols and procedures and granted permission to collect fish samples (APN project n° 1173 and S.C de Bariloche Council note n° 412/SSMA/15).

### Analysis of data

Influence of the presence of a given species on the chance of other species being present was tested using a contingency table approach, asking whether the probability of a species being present was contingent on the presence of other species. We calculated the expected counts for each unique combination of species (including basins with no species at all) assuming that the frequency of each combination is the product of the probability of each species to be present, and compared them to the observed counts using a Fisherś exact Test for contingency tables.

We then used generalized linear models (GLMs) on presence/absence and relative abundance data to assess the filtering influence of landscape and local variables. For each level and response variable type, we tried two approaches: 1) reducing the dimensionality of the independent variables through a principal components analysis, and then using the principal components explaining most variance as predictor variables in the GLM; and 2) by selecting independent variables that were not significantly correlated and using them as predictor variables in the GLM. While these two approaches are expected to yield qualitatively similar results, the second approach lends to more straightforward interpretations. All analyses were run using the function *glm* from the base package of the R computing environment R Core Team, version 3.5.1 (2018). To regress environmental variables using presence/absence as a response variable, we used a binomial logistic regression with a logit link. When the response variable was the logarithm of relative abundance, we used a Gaussian regression with an identity link.

## RESULTS

### Introduced salmonid species are unevenly distributed among basins

Only two specimens of native species were caught: one of *Galaxias maculatus* (Jennyns, 1842) in the Frey basin and another of *Hatcheria macraei* (Girard, 1885) in the Ñirihuau basin. In contrast, a total of 4531 salmonids were caught (Table C-Appendix section). We found individuals aged between 0 and 3 years old at all sites, but older specimens (up to 8 years old) were caught only in some of the Eastern basins. No fish were captured in the Newbery, Blanco, Bravo and Uhueco basins; these streams present physical barriers (large waterfalls) restricting upstream movement of fish from the rest of the system (Figure 2).

**Figure 2.**
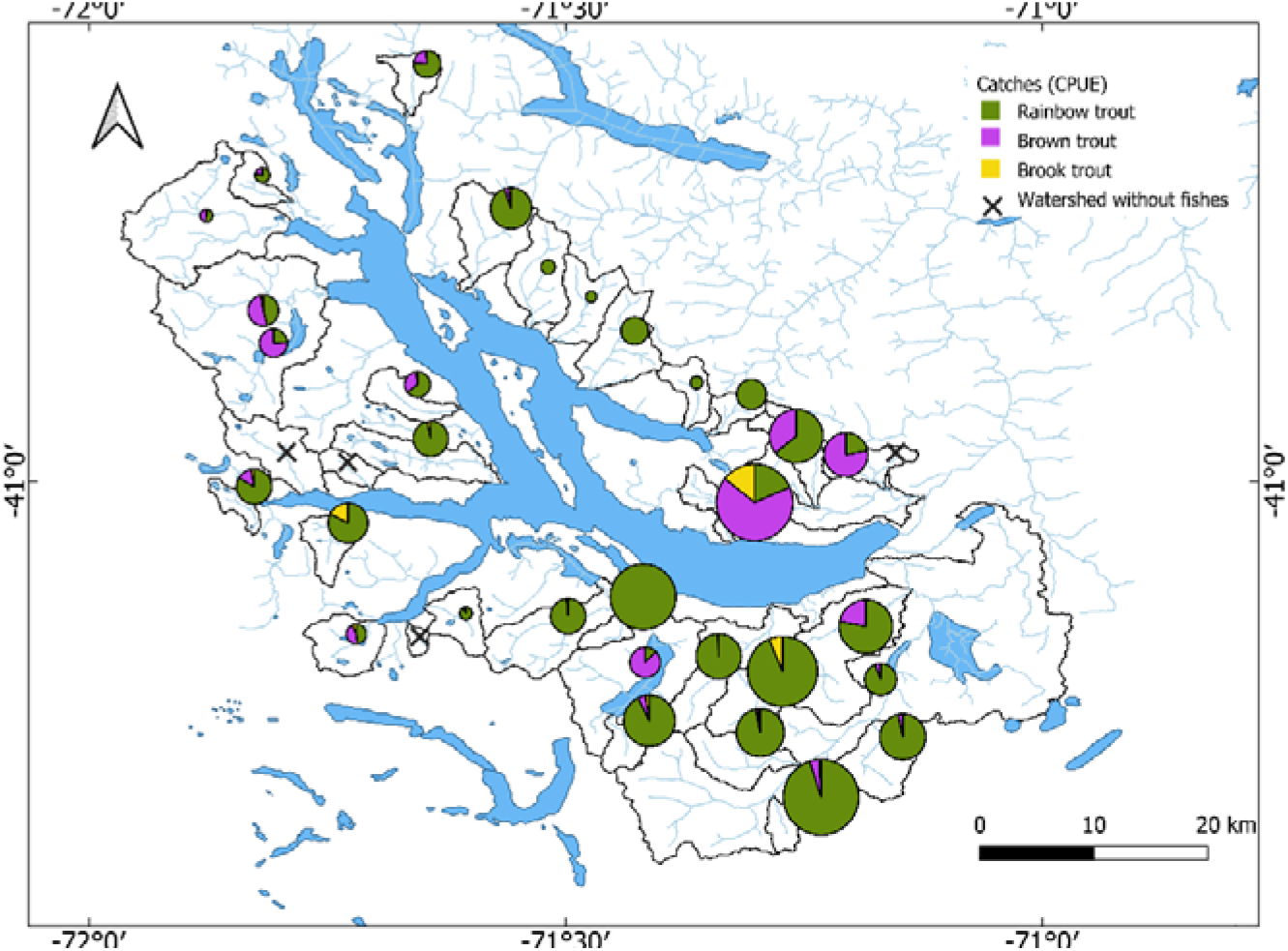
Proportion of salmonid species captured in sampled streams. The size of the pie chart is proportional to the total density of fish caught in each watershed. The crosses indicate watershed without fishes. Rainbow trout (*Oncorhynchus* mykiss), brown trout (Salmo trutta), brook trout (Salvelinus fontinalis).

*Oncorhynchus mykiss* was present in all basins where fish were caught. This species was also the most abundant in almost all basins (31/35); *Salmo trutta*, though present in 21/35 basins, was dominant in only 4 of them (Figure 2). *Salvelinus fontinalis*, was captured in 15/35 basins, but never dominated the assemblage (Figure 2).

Co-occurrence of introduced salmonid species varied among basins. The three species were found together in 10 basins. Only two out of three possible bispecific assemblages were found: S.*trutta* with O. *mykiss* (n = 10) and S. *fontinalis* with O. *mykiss* (n = 4). Monospecific assemblages were found only for O. *mykiss* (, n = 7).

### No interspecific reciprocal influences on presence are evidenced by salmonid species distribution

If the chance of a given species being present on a given basin is not influenced by the distribution of the other species (i.e., they are independent), then the chance of finding any given combination of species is simply the product of finding each species separately. We calculated the expected frequency of each combination of species under the assumption that species distributions are independent of each other, and then calculated the expected counts of all combinations using a Fisher’s test, and found no significant difference (p=0.75), suggesting that species interactions are not needed to explain the frequency of each combination (Table 1a). Expected and observed counts are even more similar when basins without fish captures are excluded from the calculations (p=0.93, Table 1b).

**Table 1.** Observed and expected frequencies for the salmonid species used for Fisherś Test **a)** including basins without caught fish and **b)** excluding basins without fishes.

| Species | Expected | Observed |
| --- | --- | --- |
| *None* | 1 | 4 |
| *Oncorhynchus mykiss* | 7 | 7 |
| *Salvelinus fontinalis* | 1 | 0 |
| *Salmo trutta* | 1 | 0 |
| *O. mykiss* x *S. fontinalis* | 5 | 4 |
| *O. mykiss* x *S. trutta* | 11 | 10 |
| *S. trutta* x *S. fontinalis* | 1 | 0 |
| *O. mykiss* x *S. fontinalis* x *S. trutta* | 8 | 10 |

**b)**
| Species | Expected | Observed |
| --- | --- | --- |
| *Oncorhynchus mykiss* | 5 | 7 |
| *Salvelinus fontinalis* | 0 | 0 |
| *Salmo trutta* | 0 | 0 |
| *O. mykiss* x *S. fontinalis* | 5 | 4 |
| *O. mykiss* x *S. trutta* | 11 | 10 |
| *S. trutta* x *S. fontinalis* | 0 | 0 |
| *O. mykiss* x *S. fontinalis* x *S. trutta* | 10 | 10 |

### Salmonid Abundances varied with environmental variables

Total catch of salmonids in streams from the upper Limay basin increased along a NW-SE gradient (Figure2). This gradient in relative abundance was associated to basins having a larger proportion of lowland, shrubby environments characterized by low rainfall, open woodlands and fewer rocky outcrops. This pattern in total catch was likely driven by dominance of O. *mykiss*, since its relative abundances presented the same pattern as the total catch (with the addition that abundance was higher at lower basins).

In contrast, *S. trutta* abundances were not clearly associated with geographic gradients (Table 2). Instead, higher abundances were associated with lower, warmer and flatter basins, with fewer High-Andean habitat and more wetlands. *Salvelinus fontinalis* abundance showed no significant correlations with any environmental variable. However, spotty occurrence and low overall abundance of this species resulted in very low statistical power to detect any existing correlations (Table 2).

**Table 2.** Spearman Rank Correlations for total and per specie catch of salmonid per unit effort (CPUEN) in relation to basin variables (n = 31). Only statistically significant differences were included in the table (p-value <0.005).

| Variables | R- Spearman | Statistic | P- values |
| --- | --- | --- | --- |
| **Salmónids CPUEN** |  |  |  |
| Latitude | -0.63 | -4.36 | 0.0001 |
| Longitude | 0.56 | 3.64 | 0.0010 |
| Rocky outcrop | 0.45 | 2.70 | 0.0115 |
| Open woodland | 0.47 | 2.90 | 0.0070 |
| Rocky summit | -0.63 | -4.34 | 0.0002 |
| Coihue Forest | -0.66 | -4.76 | 0.0001 |
| NDVI | -0.45 | -2.68 | 0.0121 |
| Precipitation | -0.59 | -3.92 | 0.0005 |
| Steppe | 0.49 | 3.01 | 0.0054 |
| ***O. mykiss* CPUEN** |  |  |  |
| Latitude | -0.61 | -4.19 | 0.0002 |
| Longitude | 0.49 | 3.06 | 0.0047 |
| Rocky outcrop | 0.45 | 2.69 | 0.0116 |
| Rocky summit | -0.44 | -2.64 | 0.0133 |
| Coihue Forest | -0.56 | -3.63 | 0.0011 |
| Precipitations | -0.48 | -2.94 | 0.0064 |
| ***S. trutta* CPUEN** |  |  |  |
| Relief | -0.42 | -2.47 | 0.0198 |
| Z max | -0.40 | -2.40 | 0.0238 |
| Rocky summit | -0.45 | -2.70 | 0.0115 |
| Mallin | 0.63 | 4.41 | 0.0001 |
| Steppe | 0.37 | 2.32 | 0.0274 |
| Wetland | 0.40 | 2.35 | 0.0256 |
| High-Andean | -0.48 | -2.93 | 0.0065 |
| Air Temperature | 0.47 | 2.83 | 0.0083 |

**Table 3.**
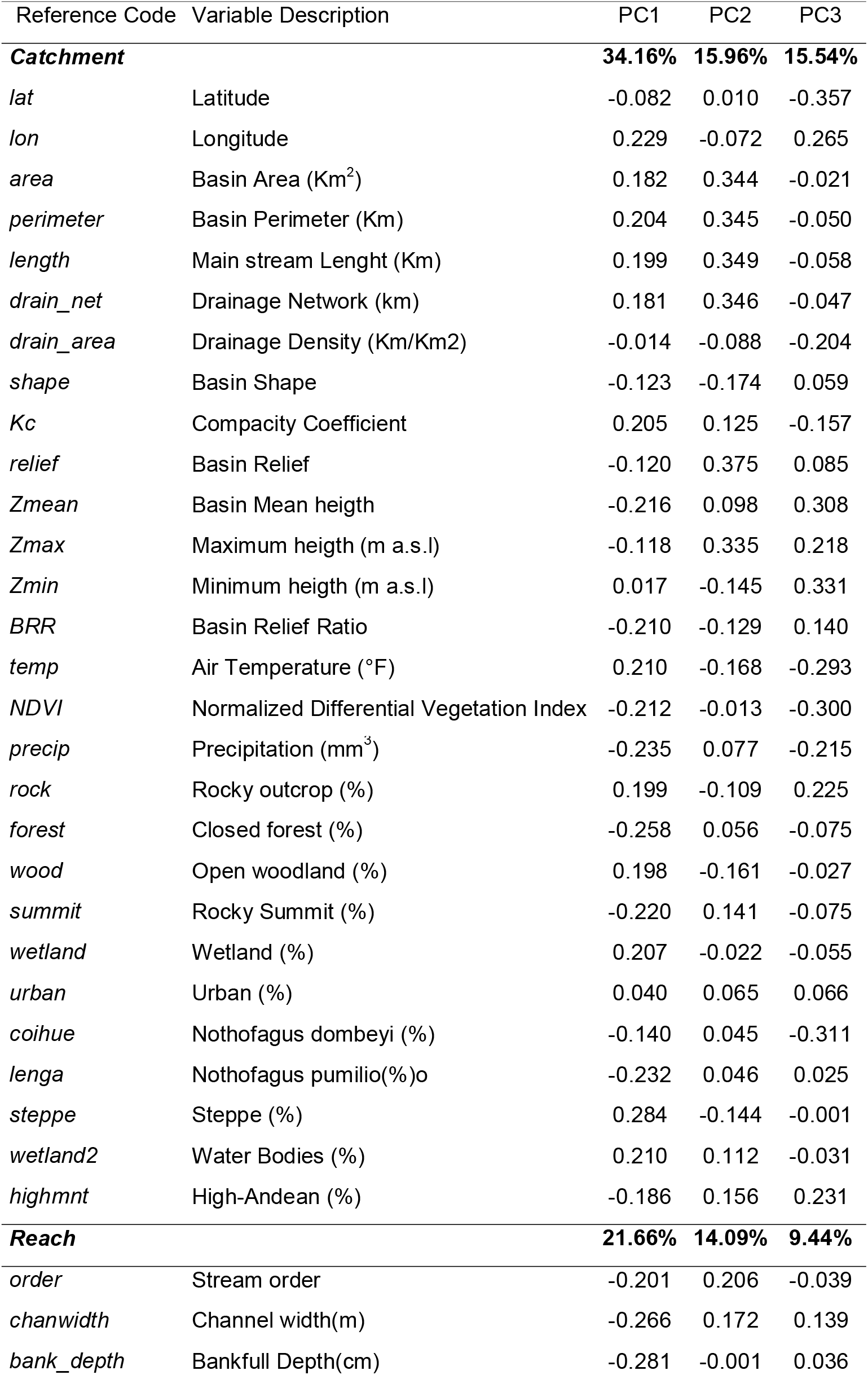
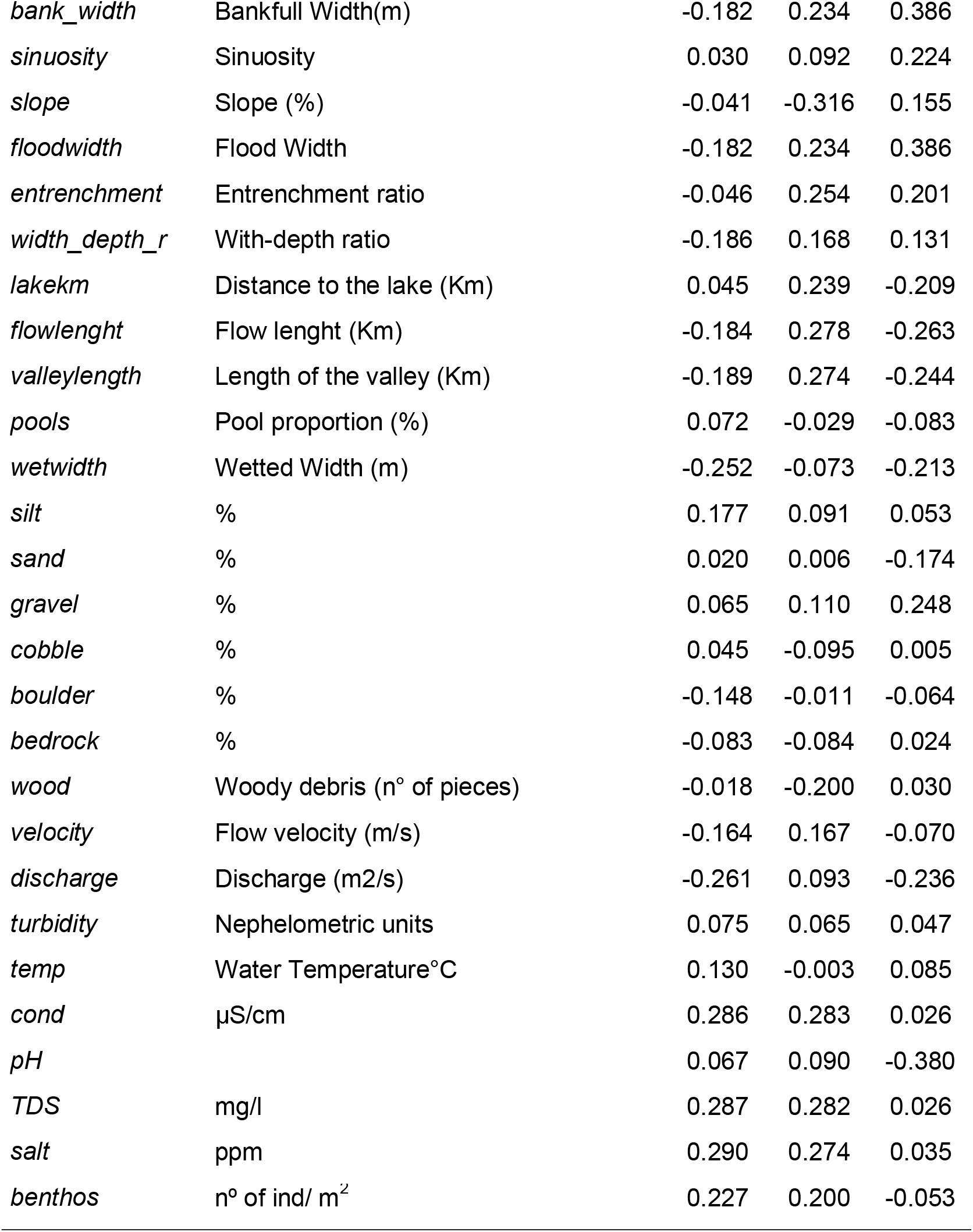
Reference codes and descriptions of habitat variables measured at two different spatial scales (catchment and reach) in the Limay river basin. Variable loadings and percent variance explained is given for the first three axes of principal component analyses to summarize variation in measurements across sites.

| Reference Code | Variable Description | PC1 | PC2 | PC3 |
| --- | --- | --- | --- | --- |
| **Catchment** |  | **34.16%** | **15.96%** | **15.54%** |
| *lat* | Latitude | -0.082 | 0.010 | -0.357 |
| *lon* | Longitude | 0.229 | -0.072 | 0.265 |
| *area* | Basin Area (Km<sup>2</sup>) | 0.182 | 0.344 | -0.021 |
| *perimeter* | Basin Perimeter (Km) | 0.204 | 0.345 | -0.050 |
| *length* | Main stream Lenght (Km) | 0.199 | 0.349 | -0.058 |
| *drain_net* | Drainage Network (km) | 0.181 | 0.346 | -0.047 |
| *drain_area* | Drainage Density (Km/Km2) | -0.014 | -0.088 | -0.204 |
| *shape* | Basin Shape | -0.123 | -0.174 | 0.059 |
| *Kc* | Compacity Coefficient | 0.205 | 0.125 | -0.157 |
| *relief* | Basin Relief | -0.120 | 0.375 | 0.085 |
| *Zmean* | Basin Mean heigth | -0.216 | 0.098 | 0.308 |
| *Zmax* | Maximum heigth (m a.s.l) | -0.118 | 0.335 | 0.218 |
| *Zmin* | Minimum heigth (m a.s.l) | 0.017 | -0.145 | 0.331 |
| *BRR* | Basin Relief Ratio | -0.210 | -0.129 | 0.140 |
| *temp* | Air Temperature (°F) | 0.210 | -0.168 | -0.293 |
| *NDVI* | Normalized Differential Vegetation Index | -0.212 | -0.013 | -0.300 |
| *precip* | Precipitation (mm<sup>3</sup>) | -0.235 | 0.077 | -0.215 |
| *rock* | Rocky outcrop (%) | 0.199 | -0.109 | 0.225 |
| *forest* | Closed forest (%) | -0.258 | 0.056 | -0.075 |
| *wood* | Open woodland (%) | 0.198 | -0.161 | -0.027 |
| *summit* | Rocky Summit (%) | -0.220 | 0.141 | -0.075 |
| *wetland* | Wetland (%) | 0.207 | -0.022 | -0.055 |
| *urban* | Urban (%) | 0.040 | 0.065 | 0.066 |
| *coihue* | Nothofagus dombeyi (%) | -0.140 | 0.045 | -0.311 |
| *lenga* | Nothofagus pumilio(%)o | -0.232 | 0.046 | 0.025 |
| *steppe* | Steppe (%) | 0.284 | -0.144 | -0.001 |
| *wetland2* | Water Bodies (%) | 0.210 | 0.112 | -0.031 |
| *highmnt* | High-Andean (%) | -0.186 | 0.156 | 0.231 |
| **Reach** |  | **21.66%** | **14.09%** | **9.44%** |
| *order* | Stream order | -0.201 | 0.206 | -0.039 |
| *chanwidth* | Channel width(m) | -0.266 | 0.172 | 0.139 |
| *bank_depth* | Bankfull Depth(cm) | -0.281 | -0.001 | 0.036 |
| Tables – Lallement, Rechencq and Zattara 2022 |  |  |  |  |
| *bank_width* | Bankfull Width(m) | -0.182 | 0.234 | 0.386 |
| *sinuosity* | Sinuosity | 0.030 | 0.092 | 0.224 |
| *slope* | Slope (%) | -0.041 | -0.316 | 0.155 |
| *floodwidth* | Flood Width | -0.182 | 0.234 | 0.386 |
| *entrenchment* | Entrenchment ratio | -0.046 | 0.254 | 0.201 |
| *width_depth_r* | With-depth ratio | -0.186 | 0.168 | 0.131 |
| *lakekm* | Distance to the lake (Km) | 0.045 | 0.239 | -0.209 |
| *flowlength* | Flow lenght (Km) | -0.184 | 0.278 | -0.263 |
| *valleylength* | Length of the valley (Km) | -0.189 | 0.274 | -0.244 |
| *pools* | Pool proportion (%) | 0.072 | -0.029 | -0.083 |
| *wetwidth* | Wetted Width (m) | -0.252 | -0.073 | -0.213 |
| *silt* | % | 0.177 | 0.091 | 0.053 |
| *sand* | % | 0.020 | 0.006 | -0.174 |
| *gravel* | % | 0.065 | 0.110 | 0.248 |
| *cobble* | % | 0.045 | -0.095 | 0.005 |
| *boulder* | % | -0.148 | -0.011 | -0.064 |
| *bedrock* | % | -0.083 | -0.084 | 0.024 |
| *wood* | Woody debris (n° of pieces) | -0.018 | -0.200 | 0.030 |
| *velocity* | Flow velocity (m/s) | -0.164 | 0.167 | -0.070 |
| *discharge* | Discharge (m2/s) | -0.261 | 0.093 | -0.236 |
| *turbidity* | Nephelometric units | 0.075 | 0.065 | 0.047 |
| *temp* | Water Temperature°C | 0.130 | -0.003 | 0.085 |
| *cond* | µS/cm | 0.286 | 0.283 | 0.026 |
| *pH* |  | 0.067 | 0.090 | -0.380 |
| *TDS* | mg/l | 0.287 | 0.282 | 0.026 |
| *salt* | ppm | 0.290 | 0.274 | 0.035 |
| *benthos* | n° of ind/ m<sup>2</sup> | 0.227 | 0.200 | -0.053 |

### Patterns of salmonid distribution are not strongly driven by landscape variables

Having ruled out a role for the interaction between species in their joint distribution (see above), we evaluated the influence of basin characteristics on individual species presence and abundances. We derived 28 watershed variables from remote sensing and topographical data sources. We found that many of the variables were highly correlated; thus, we used a principal component analysis (PCA) to reduce the dimensionality of variable space. The first three principal components (PC) explained 65.65 % of the total variance (PC1 34.16%; PC2 15.96%, PC3 15.54%).

We used these three PCs to test if basin characteristics influenced the presence of each species by fitting a logistic regression using GLMs (see Methods), but found no significant influence of either PC on presence for any of the species. We then repeated the same analysis using relative abundance (individuals caught per unit effort; CPUEN) as a dependent variable instead. We found that *O. mykiss* abundance was significantly influenced by PC3 (p=0.001, loaded mainly by Latitude and Zmin), while *S. trutta* abundance was instead, significantly influenced by PC1 (p=0.002, loaded mainly by Steppe and precipitations). *Salvelinus fontinalis* was not influenced by any PCs. Repeating the analyses using the logarithm of relative abundance as response variable yielded no significant results for any species or PCs.

Hence, for all three species, we found that the probability of a species being present in a basin was not explained by any of the basin characteristics we used in this analysis. However, we detected evidence that the abundance of *O. mykiss and S. trutta* (but not *S. fontinalis*) is influenced by different combinations of basin traits. Thus, the three species differ in their response to landscape level filtering variables.

### Local variables influence weakly but differentially the abundance of each species

To evaluate the influence of local characteristics on species presence we estimated 31 variables using local data records (see TableB-appendix section). Since several local-scale variables presented significant correlations, we used PCA to reduce the dimensionality of the variable space. The first three principal components (PC) explained 45.19 % of the total variance (PC1 21.66 %; PC2 14.09%, PC3 9.44%).

We tested whether local characteristics influenced the presence-absence of each species by fitting a logistic regression using Generalized Linear Models, and found no significant influence of either PCs on presence-absence for *O. mykiss and S. trutta*, while PC2 (loaded mainly by water quality and slope variables) had a significant influence on presence of *S. fontinalis* (z-value=-2.62, p-value=0.009). We then tested if reach variables influence relative abundances (either absolute values or their logarithms) and found a significant relation between PC1 (loaded mainly by channel morphology and water quality variables) and *O. mykiss* abundances (absolute – t=3.86, p=0.0004 and logarithms values –t=2.94, p=0.0055), but no significant influence on *S. trutta*. We found significant influence of PC2 on the logarithm of *S. fontinalis* abundance (t-value=-2.97, p=0.0052) but not on the absolute values. In summary, our regression analysis showed that the influence of local traits varies across all three salmonid species.

## DISCUSSION

One of the most pervasive concepts in the study of community assembly is the metaphor of the environmental filter, which refers to abiotic factors that prevent the establishment or persistence of species in a particular location. However, this concept has been criticized because the evidence used in many studies to assess environmental filtering is insufficient to distinguish filtering from the evolutionary outcome of historical biotic interactions (Kraft *et al*., 2015). In our work we took advantage of the relatively recent and well-studied history of salmonid introductions to evaluate if non-native species show different patterns of association with abiotic factors at different spatial scales of the filter.

We found widespread presence of salmonid species and an almost total absence of native species in our study area. Thus, the only expected source of current biotic interference would be interspecific interaction between the salmonid species themselves. However, the results of our contingence analysis on species distribution does not support a scenario in which any of the species is affecting the distribution of the other species. This suggests that streams in our study area have not reached carrying capacity for salmonids and that they might be partitioning the riverine habitats to minimize niche overlap, as reported in other parts of the world (Bozek & Hubert, 1992; Reeves, Bisson, & Dambacher 1998; Fausch, 2008; Marchetti *et al*., 2011).

Interspecific interference among salmonid species has been proposed in previous studies within this region (Juncos, Beauchamp & Vigliano, 2013; Arismendi *et al*., 2014); we found however no evidence for it. Thus, we can assume that interference has played only a minor role in determining current fish distribution in the region. Instead, current patterns of presence and abundance of salmonids are best explained as the product of environmental filters. This agrees with studies relating the success of salmonid invasions in southern Chile to the excellent abiotic conditions they found in the region (Pascual *et al*., 2007; Habit *et al*., 2012, 2015).

Biological invasions are inherently complex. A successful invader must survive a series of events: transport to the invasion site, initial establishment, spread to a broad area, and then integration into the existing biotic community (Moyle & Light, 1996; Kolar & Lodge, 2001). Not surprisingly, most introduced species fail to become established and never reach invasive status (Moyle & Light, 1996; Arismendi *et al*., 2014). Our results suggest that in North Patagonia, biotic resistance from native fish species seems to have had little or no influence on the invasion process by introduced salmonid species, and that success of invaders in the face of low odds is related, as we previously suggested, mostly to the presence of favorable environments, such as adequate flow regimes (conditioned for the rainfall gradient) and food availability (Lallement *et al*., 2020).

In contrast to other reports (Marchetti, Moyle & Levine, 2004; Stanfield, Scott & Borwick, 2006b), we saw no evidence that presence/absence patterns of salmonid distribution were strongly driven by landscape variables, except for those basins with environmental barriers. However, when analyzing responses of relative abundances (CPUEN), the influence of climatic and geomorphological variables (e.g., precipitation and relief) became more evident. These two types of factors have been mentioned in other systems as determinants of salmonid distribution (Stanfield *et al*., 2006b; Warren, Dunbar & Smith, 2015): rainfall and geology have a direct influence on stream discharge and are thus determinant during early development of salmonid life cycles (Heggenes & Traaen, 1988; Nehring & Anderson, 1993). Determining the relative influence of landscape and habitat features seems likely to remain a difficult problem because their influences may be inextricably linked in most situations.

Landscape characteristics (e.g., general slope of the valley and geomorphological aspects) determine local river section traits such as substrate composition, pool dimensions and refuge availability, which in turn strongly correlate with the structure and distribution of the assemblages of fish in a basin (Fischer & Paukert, 2008; Rowe, Pierce & Wilton, 2009). In our study area, we found evidence that some local traits modulate O. *mykiss* abundance but do not explain abundances of S. *trutta* and *S. fontinalis*, or the presence-absence patterns for all three salmonid species. Low abundances for a species could be causing diminished statistical power to detect the influence of environmental variables; this could be affecting our results for brook trout. Differential abundances are nonetheless likely to result from differential responses to the same environment by each species. Thus, the layered influence of environmental filters was reflected in a weak but differential influence of local traits on abundance of each salmonid species.

Habitat suitability for salmonids is often controlled by local variables, such as temperature regime (Rahel & Nibbelink, 1999; Harig & Fausch, 2002; Coleman & Fausch, 2007), flow regime (Strange & Foin, 1999; Fausch *et al*., 2001), stream size (Rahel & Nibbelink, 1999) and habitat factors correlated with stream gradient and channel geomorphology (Fausch, 1989; Montgomery *et al*., 1999). Due to the steep precipitation gradient in our study area, several streams originate in wooded areas and cross wide valleys where shrubby vegetation typical of arid steppes predominates. Shrubby riverbanks result in higher solar irradiation and air temperatures, favoring higher primary production of the periphyton and sustaining an important biomass of macroinvertebrates (Miserendino, 2007; Modenutti *et al*., 2010). It is within these streams where we found higher fish abundances.

Our results are interesting in light of other studies that have tried to identify the relative importance of different process scales in structuring biotic assemblages. Previous predictive modeling studies have indicated that landscape-scale watershed characteristics explain better fish distributions than reach-scale characteristics (Creque, Rutherford & Zorn, 2005; Frimpong *et al*., 2005). However, our study did not find that watershed-scale variables were significantly better at explaining or predicting fish distributions in North Patagonian streams. This could be due to a lack of habitat saturation by fishes resulting from temporal variability in fish distributions, to us failing to measure the reach-and watershed-scale variables actually driving distribution, or to an insufficient sample size. As Stanfield et al. (2006a), we do not contend that landscape variables are the most important for salmonids, but suggest that landscape traits do condition the range of geomorphological variation inside a watershed that ultimately defines the present local abundance at the reach level (Lallement *et al*., 2020). In any case, further information is still needed to fully understand how the variability of spatial scale affects the distribution of fishes in streams of Patagonia.

Even though restoring streams to the conditions prior to the introductions is not feasible due to the wide distribution and dominance of salmonids at present, a possible strategy for conservation would be to focus the efforts on particular environments still devoid of salmonids. The existence of barriers that prevent salmonid colonization of headwaters has been shown to lead to different biota assemblages (Buria *et al*., 2007; Albariño & Buria, 2011). Given our results, a possible strategy would be to determine the effect of differential exclusions on the stream biota, as well as the effect of such exclusions measured on the general pool of salmonids. Thus, if it is found that the fishing stock depends on sourcing from a few basins, management efforts could concentrate in those sources.

Although not explored in this study, distributions of fishes are variable and may be regulated by seasonal movements and the frequency and duration of stochastic flow cycles. In any case, it is important to bear in mind that the results obtained in regions with different histories of colonization and introduction are not necessarily extrapolable. From a management perspective, our results suggest that considering the pristine (or near-pristine) condition of the streams sampled here, the relationships observed between fishes and landscape variables can be used, for example, as a reference for further studies addressing the effects of human modifications on aquatic biodiversity of North Patagonia. We believe that our findings expand our current understanding about how the combination of factors (habitat template) across different scales structure aquatic assemblages within Patagonia and can be used to strategically guide future management actions.

## Supporting information

Appendix Section (Tables A,AI,AII,AIII, B and C)

## Acknowledgments

We thank Alejandro Sosnovsky and Patricio Macchi for their comments on an earlier draft, and to the National Park Administration (APN) that through the Biodiversity Information System (SIB) provided us with the data for Use of Land and Vegetation in watersheds. We thank all the members of the Grupo de Evaluación y Manejo de Recursos Icticos (GEMaRI) for field assistance. Funding was provided by the Agencia Nacional de Promoción de Ciencia y Tecnología, Argentina (PICT projects 016 and 2959). The authors declare no conflict of interest.

## Data Availability Statement

The data and code with the analyses supporting our findings, along with a fully explicit RMarkdown version of this manuscript and its references, are available at https://github.com/ezattara/patagonian_salmonid_distribution

## AUTHORS’ CONTRIBUTIONS

Conceptualization: MEL, EEZ; data curation: MEL, EEZ; formal analysis: MEL, EEZ; investigation: MEL, EEZ, MR; visualization: MEL, MR; writing – original draft: MEL, EEZ; writing – review & editing: MEL, EEZ, MR.

## Notes

Instituto de Matemática Aplicada San Luis, IMASL, CONICET and Universidad Nacional de San Luis, Ejército de los Andes 950, D5700HHW San Luis, Argentina

https://github.com/ezattara/patagonian_salmonid_distribution

### Competing Interest Statement

The authors have declared no competing interest.

