## Appendix Section (Tables A,AI,AII,AIII, B and C) for "The Devil Is In The Details: Landscape Features Are Insufficient To Explain Patterns Of Non-Native Fishes Distribution In North Patagonian Streams"

apA

**Table A** Group of basin variables measured with Qgis. Reference code column correspond to the variables names used in the R script.

| Geomorphology | Reference Code | Variable Description |
|---|---|---|
| Basin Area | *lat* | BA ($Km^2$) |
| Basin Perimeter | *lon* | BP (Km) |
| Main Stream Length | *length* | MSL (Km) |
| Drainage Network | *drain_net* | DN=Σ stream length (Km) |
| Drainage Density | *drain_area* | DD=RD/A ($Km/Km^2$) |
| Basin Shape | *shape* | BS=DD/(main channel length)$^2$ |
| Compacity Coefficient | *Kc* | Kc=0.28(P/√A) |
| Basin Relief | *Relief* | BR=Highest basin point – Lower basin point |
| Basin Relief Ratio | *BRR* | BRR=RC – stream length |
| Maximum heigth | *Zmax* | Zmax= Highest basin point (m.a.s.l) |
| Average heigth | *Zmean* | Zav= Average basin height (m.a.s.l) |
| Minimum heigth | *Zmin* | Zmin=Lower basin point (m.a.s.l) |
| Land Use | | |
| Rocky outcrop | *Rock* | |
| Gravel | *Gra* | |
| Closed Forest | *forest* | % of basin occupied with the specific category |
| Open woodlands | *wood* | |
| Rocky summit | *summit* | |
| Water Bodies (WB) | *wetland2* | |
| Urban | *urban* | |
| Plantation | | |
| Clearence | | |
| Vegetation | | |
| *Fitzroya cupressoides* | | (Alerce) |
| *Austrocedrus chilensis* | | (Ciprés de la cordillera) |
| *Nothofagus dombeyi* | *coihue* | (Coihue) |
| *Nothofagus pumilio* | *lenga* | (Lenga) |
| *Nothofagus antárctica* | | (Ñire) |
| Steepe | *steppe* | |
| Wetland | *wetland* | |
| High-Andean | *highmnt* | |
| Climatological | | |
| Summer Mean Temperature | *Temp* | T (°F) |
| Mean Annual Precipitation | *precip* | Precipitation ($mm^3$) |

### Appendix

apA

| | | |
|---|---|---|
| Normalized Differential Vegetation Index | *NDVI* | NDVI |

apA

**Table A-I** Basin geomorphology variables calculated for the 35 watersheds.

| Basin name | Latitude | Longitude | BA | BP | MSL | DN | DD | BS | Kc | BRR | BR | Elevation | Z av | Zmax | Zmin |
|---|---|---|---|---|---|---|---|---|---|---|---|---|---|---|---|
| **2.Acantuco** | -40.6870 | -71.8259 | 18.76 | 21.67 | 6.64 | 8.80 | 0.47 | 0.0106 | 1.40 | 175.34 | 1165.00 | 1351.50 | 1286.18 | 1934.00 | 769.00 |
| **8.Blanco** | -40.9863 | -71.7269 | 14.52 | 17.75 | 4.72 | 4.72 | 0.32 | 0.0146 | 1.30 | 263.11 | 1241.00 | 1393.00 | 1410.95 | 2014.00 | 773.00 |
| **10.Blest** | -41.0244 | -71.8452 | 12.5 | 20.48 | 6.80 | 10.39 | 0.83 | 0.0180 | 1.62 | 151.63 | 1031.00 | 1281.50 | 1227.72 | 1797.00 | 766.00 |
| **15.Bonito** | -40.7357 | -71.5788 | 56.72 | 39.23 | 13.24 | 22.74 | 0.40 | 0.0023 | 1.46 | 86.65 | 1147.00 | 1342.50 | 1384.99 | 1916.00 | 769.00 |
| **7.Bravo** | -40.9687 | -71.8035 | 46.91 | 38.84 | 10.38 | 12.61 | 0.27 | 0.0025 | 1.59 | 109.08 | 1132.00 | 1333.00 | 1263.73 | 1901.00 | 769.00 |
| **25.Casa de Piedra** | -41.1604 | -71.5157 | 64.78 | 44.05 | 19.77 | 34.13 | 0.53 | 0.0013 | 1.53 | 73.70 | 1457.00 | 1477.50 | 1473.44 | 2206.00 | 749.00 |
| **26.Cascada** | -41.1561 | -71.4530 | 12.53 | 20.42 | 7.67 | 7.67 | 0.61 | 0.0104 | 1.62 | 176.58 | 1355.00 | 1487.50 | 1256.49 | 2149.00 | 794.00 |
| **24.Castilla** | -41.0226 | -71.3418 | 25.76 | 31 | 10.27 | 21.49 | 0.83 | 0.0079 | 1.71 | 63.67 | 654.00 | 1099.00 | 925.18 | 1426.00 | 772.00 |
| **23.Chacabuco** | -40.9954 | -71.2287 | 134.94 | 70.89 | 25.73 | 68.23 | 0.51 | 0.0008 | 1.71 | 46.20 | 1189.00 | 1359.00 | 1054.93 | 1954.00 | 765.00 |
| **32.Challhuaco** | -41.2366 | -71.3091 | 41.62 | 28.95 | 9.56 | 12.92 | 0.31 | 0.0034 | 1.26 | 134.85 | 1289.00 | 1779.50 | 1450.07 | 2228.00 | 939.00 |
| **6.Coluco** | -40.9123 | -71.6688 | 25.15 | 25.93 | 9.26 | 9.26 | 0.37 | 0.0043 | 1.45 | 127.71 | 1182.00 | 1362.00 | 1438.06 | 1953.00 | 771.00 |
| **35.De la Quebrada** | -41.3616 | -71.2715 | 14.31 | 18.53 | 5.22 | 5.22 | 0.36 | 0.0134 | 1.37 | 185.57 | 969.00 | 2000.50 | 1635.22 | 2103.00 | 1134.00 |
| **29.Del Medio** | -41.1808 | -71.2129 | 108.07 | 60.25 | 24.52 | 35.20 | 0.33 | 0.0005 | 1.62 | 38.14 | 935.00 | 1217.00 | 979.40 | 1727.00 | 792.00 |
| **16.Estacada** | -40.7830 | -71.5257 | 49.06 | 35.55 | 13.87 | 27.06 | 0.55 | 0.0029 | 1.42 | 86.87 | 1205.00 | 1373.50 | 1447.21 | 1976.00 | 771.00 |
| **12.Frey** | -41.1712 | -71.7300 | 36.59 | 26.23 | 7.49 | 9.84 | 0.27 | 0.0048 | 1.21 | 185.76 | 1391.00 | 1465.50 | 1363.49 | 2159.00 | 768.00 |
| **5.Gallardo** | -40.8701 | -71.8212 | 97.32 | 52.24 | 17.38 | 32.01 | 0.33 | 0.0011 | 1.48 | 66.76 | 1160.00 | 1362.00 | 1273.73 | 1942.00 | 782.00 |
| **27.Gutierrez** | -41.2067 | -71.4326 | 160.17 | 63.17 | 28.73 | 60.57 | 0.38 | 0.0005 | 1.40 | 56.18 | 1614.00 | 1576.00 | 1261.34 | 2383.00 | 769.00 |
| **18.Huemul** | -40.8569 | -71.4419 | 54.14 | 36.52 | 12.80 | 34.12 | 0.63 | 0.0038 | 1.39 | 107.09 | 1371.00 | 1446.50 | 1416.06 | 2140.00 | 769.00 |
| **34.Las Minas** | -41.2928 | -71.1703 | 44.4 | 31.58 | 10.59 | 15.21 | 0.34 | 0.0031 | 1.33 | 53.14 | 563.00 | 1322.50 | 1190.42 | 1503.00 | 940.00 |
| **14.LLuvuco** | -41.1457 | -71.6111 | 27.19 | 24.98 | 6.40 | 9.01 | 0.33 | 0.0081 | 1.34 | 216.94 | 1389.00 | 1497.00 | 1537.49 | 2190.00 | 801.00 |
| **4.Machete** | -40.8373 | -71.8332 | 193.67 | 72.96 | 26.40 | 88.88 | 0.46 | 0.0007 | 1.47 | 45.07 | 1190.00 | 1350.00 | 1279.72 | 1945.00 | 755.00 |
| **21.Manzano-Jones** | -40.9812 | -71.2790 | 30.49 | 37.4 | 13.10 | 13.38 | 0.44 | 0.0026 | 1.90 | 79.23 | 1038.00 | 1336.00 | 1174.19 | 1892.00 | 854.00 |
| **9.Millaqueo** | -40.9743 | -71.6598 | 52 | 42.31 | 16.96 | 35.64 | 0.69 | 0.0024 | 1.64 | 73.81 | 1252.00 | 1392.00 | 1353.28 | 2018.00 | 766.00 |
| **1.Neuquenco** | -40.5768 | -71.6595 | 21.94 | 26.31 | 6.96 | 7.42 | 0.34 | 0.0070 | 1.57 | 156.24 | 1088.00 | 1323.00 | 1128.61 | 1871.00 | 783.00 |

**Table A-I** Continuation

| Basin name | Latitude | Longitude | BA | BP | MSL | DN | DD | BS | Kc | BRR | BR | Elevation | Z av | Zmax | Zmin |
|---|---|---|---|---|---|---|---|---|---|---|---|---|---|---|---|
| **22.Newbery** | -40.9801 | -71.1867 | 27.83 | 29.27 | 6.56 | 12.97 | 0.47 | 0.0108 | 1.55 | 100.62 | 660.00 | 1183.50 | 1066.44 | 1454.00 | 794.00 |
| **28.Ñireco** | -41.2078 | -71.3215 | 113.16 | 62.83 | 19.74 | 37.55 | 0.33 | 0.0009 | 1.65 | 73.69 | 1455.00 | 1500.50 | 1292.83 | 2228.00 | 773.00 |
| **33.Ñirihuau** | -41.2253 | -71.1863 | 723.8 | 193.38 | 58.07 | 278.58 | 0.38 | 0.0001 | 2.01 | 25.30 | 1469.00 | 1499.50 | 1191.72 | 2234.00 | 765.00 |
| **11.Patiruco** | -41.0652 | -71.7491 | 24.43 | 24.84 | 6.53 | 13.00 | 0.53 | 0.0125 | 1.41 | 187.55 | 1224.00 | 1392.50 | 1271.24 | 2001.00 | 777.00 |
| **19.Pedregoso** | -40.9034 | -71.3690 | 20.53 | 25.35 | 7.22 | 8.52 | 0.41 | 0.0080 | 1.57 | 195.93 | 1415.00 | 1478.50 | 1567.94 | 2186.00 | 771.00 |
| **3.Pireco** | -40.7283 | -71.8834 | 125.48 | 61.19 | 21.53 | 59.93 | 0.48 | 0.0010 | 1.53 | 54.94 | 1183.00 | 1345.50 | 1258.26 | 1937.00 | 754.00 |
| **20.Quintriqueu** | -40.9248 | -71.3206 | 15.26 | 20.8 | 5.47 | 5.48 | 0.36 | 0.0120 | 1.49 | 209.61 | 1147.00 | 1358.00 | 1526.57 | 1932.00 | 785.00 |
| **17.Ragintuco** | -40.8126 | -71.4787 | 38.76 | 34.24 | 11.62 | 22.04 | 0.57 | 0.0042 | 1.54 | 112.25 | 1304.00 | 1423.00 | 1460.31 | 2075.00 | 771.00 |
| **30.Torrontegui** | -41.2788 | -71.4390 | 17.47 | 20.03 | 6.16 | 6.16 | 0.35 | 0.0093 | 1.34 | 217.84 | 1342.00 | 1980.00 | 1622.19 | 2145.00 | 803.00 |
| **31.Tristeza** | -41.2894 | -71.3209 | 41.29 | 32.45 | 12.16 | 15.88 | 0.38 | 0.0026 | 1.41 | 95.75 | 1164.00 | 1899.50 | 1648.61 | 2234.00 | 1070.00 |
| **13.Uhueco** | -41.1671 | -71.6557 | 5.44 | 10.82 | 3.42 | 3.42 | 0.63 | 0.0537 | 1.30 | 354.38 | 1212.00 | 1427.00 | 1530.51 | 2024.00 | 812.00 |

### Appendix

apA

**Table A-II** Land use in streams basins. Data expressed in % of the watershed occupied for each land use.

| Basin name | Rocky outcrop | Gravel | Closed forest | Open woodland | Rocky summit | Water body | Mallin | Urban | Plantation | Clearence |
|---|---|---|---|---|---|---|---|---|---|---|
| **2.Acantuco** | 0 | 0.66 | 70.76 | 0 | 28.56 | 0 | 0 | 0 | 0 | 0 |
| **8.Blanco** | 0 | 0 | 43.92 | 0 | 54.10 | 1.98 | 0 | 0 | 0 | 0 |
| **10.Blest** | 0 | 0 | 77.32 | 0 | 17.29 | 5.35 | 0 | 0 | 0 | 0 |
| **15.Bonito** | 0 | 0 | 73.58 | 0 | 26.42 | 0 | 0 | 0 | 0 | 0 |
| **7.Bravo** | 0 | 0 | 68.04 | 0 | 31.95 | 0.02 | 0 | 0 | 0 | 0 |
| **25.Casa de Piedra** | 0 | 0 | 58.11 | 10.50 | 30.75 | 0.64 | 0 | 0 | 0 | 0 |
| **26.Cascada** | 0 | 0 | 54.93 | 20.67 | 24.37 | 0 | 0 | 0 | 0 | 0 |
| **24.Castilla** | 0 | 0 | 52.79 | 47.02 | 0 | 0 | 0.209 | 0 | 0 | 0 |
| **23.Chacabuco** | 19.55 | 7.40 | 33.13 | 22.48 | 0 | 0 | 17.45 | 0 | 0 | 0 |
| **32.Challhuaco** | 20.04 | 0 | 54.40 | 21.40 | 4.16 | 0 | 0 | 0 | 0 | 0 |
| **6.Coluco** | 0 | 0 | 60.19 | 0 | 39.50 | 0.29 | 0 | 0 | 0 | 0 |
| **35.De la Quebrada** | 39.39 | 0 | 59.59 | 0 | 0 | 0 | 1.00 | 0 | 0 | 0 |
| **29.Del Medio** | 0 | 64.63 | 0 | 25.64 | 1.91 | 0 | 1.00 | 0 | 0 | 0 |
| **16.Estacada** | 0 | 0 | 67.66 | 0 | 32.33 | 0 | 7.82 | 0.10 | 0 | 0 |
| **12.Frey** | 0 | 0 | 74.71 | 0 | 21.06 | 4.23 | 0 | 0 | 0 | 0 |
| **5.Gallardo** | 0 | 0 | 63.91 | 0 | 26.78 | 8.61 | 0.704 | 0 | 0 | 0 |
| **27.Gutiérrez** | 0 | 0 | 60.22 | 10.85 | 17.55 | 11.31 | 0 | 0 | 0 | 0 |
| **18.Huemul** | 0 | 0 | 64.29 | 0 | 35.72 | 0 | 0 | 0 | 0 | 0 |
| **34.Las Minas** | 53.93 | 0 | 17.67 | 28.10 | 0 | 0 | 0.31 | 0 | 0 | 0 |
| **14.LLuvuco** | 0 | 0 | 71.01 | 0 | 27.95 | 1.04 | 0 | 0 | 0 | 0 |
| **4.Machete** | 0 | 0 | 63.42 | 0 | 28.57 | 4.78 | 3.23 | 0 | 0 | 0 |
| **21.Manzano-Jones** | 7.37 | 0 | 77.96 | 4.13 | 0 | 0 | 10.55 | 0 | 0 | 0 |
| **9.Millaqueo** | 0 | 0 | 63.32 | 0 | 36.67 | 0.01 | 0 | 0 | 0 | 0 |
| **1.Neuquenco** | 0 | 0 | 53.04 | 36.24 | 10.73 | 0 | 0 | 0 | 0 | 0 |

**Table A-II** Continuation

| Basin name | Rocky outcrop | Gravel | Closed forest | Open woodland | Rocky summit | Water body | Mallin | Urban | Plantation | Clearence |
|---|---|---|---|---|---|---|---|---|---|---|
| **22.Newbery** | 60.25 | 0.54 | 0 | 19.38 | 0 | 0 | 19.84 | 0 | 0 | 0 |
| **28.Ñireco** | 11.90 | 0 | 37.37 | 38.55 | 8.73 | 0.04 | 0 | 1.58 | 0.80 | 1.02 |
| **33.Ñirihuau** | 41.89 | 15.72 | 18.99 | 9.30 | 3.60 | 0.01 | 8.94 | 0.02 | 0.41 | 0 |
| **11.Patiruco** | 0 | 0 | 80.76 | 0 | 18.38 | 0.87 | 0 | 0 | 0 | 0 |
| **19.Pedregoso** | 0 | 0 | 45.62 | 7.32 | 47.06 | 0 | 0 | 0 | 0 | 0 |
| **3.Pireco** | 0 | 2.38 | 65.65 | 0 | 21.35 | 0.96 | 3.83 | 0 | 0 | 0 |
| **20.Quintriqueuco** | 23;17 | 0 | 68.78 | 5.04 | 3.03 | 0 | 0 | 0 | 0 | 0 |
| **17.Ragintuco** | 0 | 0 | 57.67 | 0 | 42.34 | 0 | 0 | 0 | 0 | 0 |
| **30.Torrontegui** | 0 | 0 | 61.19 | 0 | 38.27 | 0.55 | 0 | 0 | 0 | 0 |
| **31.Tristeza** | 32.36 | 0 | 63.47 | 4.19 | 0 | 0 | 0 | 0 | 0 | 0 |
| **13.Uhueco** | 0 | 0 | 70.47 | 0 | 29.16 | 0.42 | 0 | 0 | 0 | 0 |

**Table A-III** Vegetation strata (data expressed in %) and climatic variables in the sub-basins of the Limay river basins.

| | Vegetation variables | | | | | | | | Climatological variables | | |
|---|---|---|---|---|---|---|---|---|---|---|---|
| Basin name | Alerce | Ciprés cordillera | Coihue | Lenga | Ñire | Steppe | Wetland | High-Andean | Summer Temperature | NDVI | Precipitation |
| **2.Acantuco** | 0 | 0 | 27.56 | 43.02 | 0.80 | 0 | 0 | 28.62 | 53.94 | 0.22 | 2698.29 |
| **8.Blanco** | 0 | 0 | 13.71 | 40.63 | 0 | 0 | 0 | 45.66 | 48.98 | 0.11 | 2386.56 |
| **10.Blest** | 11.28 | 0 | 14.08 | 62.48 | 0 | 0 | 0 | 0 | 56.30 | 0.26 | 3113.83 |
| **15.Bonito** | 0 | 0 | 20.38 | 38.10 | 12.96 | 0 | 0 | 28.56 | 50.77 | 0.22 | 2001.43 |
| **7.Bravo** | 32.83 | 0 | 0 | 30.31 | 0 | 0 | 0 | 35.96 | 53.59 | 0.12 | 2775.10 |
| **25.Casa de** | 0 | 0 | 0.05 | 35.86 | 23.37 | 0 | 0 | 37.17 | 41.99 | 0.07 | 2324.05 |
| **26.Cascada** | 0 | 0 | 0 | 17.56 | 47.57 | 0 | 0 | 20.43 | 61.77 | 0.07 | 1926.90 |
| **24.Castilla** | 0 | 16.11 | 0 | 8.85 | 43.59 | 31.37 | 0 | 0 | 75.16 | 0.11 | 1439.01 |
| **23.Chacabuco** | 0 | 0 | 0 | 13.35 | 0 | 71.03 | 11.17 | 4.45 | 65.27 | -0.02 | 1196.57 |
| **32.Challhuaco** | 0 | 0 | 0 | 51.83 | 16.75 | 0 | 0 | 31.43 | 45.33 | 0.06 | 1834.49 |
| **6.Coluco** | 0 | 0 | 21.11 | 38.93 | 0 | 0 | 0 | 39.72 | 51.98 | 0.13 | 2541.63 |
| **35.De la** | 0 | 0 | 0 | 48.36 | 6.08 | 0 | 0 | 45.56 | 36.31 | 0.04 | 1526.00 |
| **29.Del Medio** | 0 | 0 | 0 | 7.07 | 15.00 | 72.01 | 4.52 | 1.41 | 68.12 | -0.05 | 1248.79 |
| **16.Estacada** | 0 | 0 | 22.85 | 41.70 | 0 | 0 | 0 | 35.45 | 48.62 | 0.18 | 1978.71 |
| **12.Frey** | 0 | 0 | 24.76 | 53.79 | 0 | 0 | 0 | 15.63 | 49.46 | 0.20 | 2626.64 |
| **5.Gallardo** | 0 | 0 | 17.70 | 45.91 | 0 | 0 | 0 | 27.51 | 50.81 | 0.10 | 2962.76 |
| **27.Gutiérrez** | 0 | 10.41 | 6.07 | 25.99 | 18.51 | 0 | 0 | 20.90 | 52.12 | 0.03 | 1741.80 |
| **18.Huemul** | 0 | 0 | 26.21 | 29.63 | 3.68 | 0 | 0 | 40.49 | 44.65 | 0.12 | 2112.66 |
| **34.Las Minas** | 0 | 0 | 0 | 6.58 | 0 | 93.42 | 0 | 0 | 62.07 | -0.04 | 1382.30 |
| **14.LLuvuco** | 0 | 0 | 2.98 | 51.82 | 0 | 0 | 0 | 44.21 | 40.86 | 0.04 | 2507.22 |
| **4.Machete** | 0 | 0 | 19.26 | 43.64 | 0 | 0 | 0 | 32.51 | 52.89 | 0.13 | 2942.88 |
| **21.Manzano-** | 0 | 0 | 0 | 41.52 | 0 | 49.49 | 2.33 | 6.69 | 62.25 | 0.10 | 1426.93 |
| **9.Millaqueo** | 0 | 0 | 16.13 | 30.13 | 15.88 | 0 | 0 | 35.83 | 52.01 | 0.13 | 2310.23 |
| **1.Neuquenco** | 0 | 0 | 46.17 | 21.10 | 27.21 | 0 | 0 | 5.52 | 62.26 | 0.30 | 2302.93 |

**Table A-III** Continuation

| Basin name | Vegetation variables | | | | | | | | Climatological variables | | |
|---|---|---|---|---|---|---|---|---|---|---|---|
| | Alerce | Ciprés cordillera | Coihue | Lenga | Ñire | Steppe | Wetland | High-Andean | Summer Temperature | NDVI | Precipitation |
| **22.Newbery** | 0 | 0 | 0 | 0 | 0 | 100 | 0 | 0 | 70.76 | -0.09 | 1137.50 |
| **28.Ñireco** | 0 | 0 | 0 | 30.10 | 20.46 | 5.34 | 0 | 26.90 | 51.12 | 0.01 | 1657.58 |
| **33.Ñirihuau** | 0 | 0 | 0 | 16.92 | 3.96 | 60.04 | 5.48 | 13.08 | 56.12 | -0.04 | 1203.16 |
| **11.Patiruco** | 0 | 0 | 23.82 | 58.37 | 0 | 0 | 0 | 17.81 | 56.32 | 0.17 | 2165.42 |
| **19.Pedregoso** | 0 | 5.36 | 0 | 2.19 | 7.26 | 0 | 0 | 85.24 | 39.16 | 0 | 1796.36 |
| **3.Pireco** | 0 | 0 | 26.24 | 41.35 | 2.66 | 0 | 0 | 28.80 | 54.22 | 0.19 | 2800.08 |
| **20.Quintriqueuco** | 0 | 5.57 | 0 | 55.37 | 0 | 0 | 0 | 39.06 | 43.96 | 0.12 | 1576.86 |
| **17.Ragintuco** | 0 | 0 | 14.40 | 41.51 | 6.91 | 0 | 0 | 37.18 | 46.61 | 0.15 | 2132.30 |
| **30.Torrontegui** | 0 | 6.18 | 0 | 39.38 | 6.81 | 0 | 0 | 47.17 | 36.65 | 0.03 | 1462.27 |
| **31.Tristeza** | 0 | 0 | 0 | 37.54 | 9.01 | 0.48 | 0 | 52.97 | 38.22 | -0.01 | 1733.11 |
| **13.Uhueco** | 0 | 0 | 1.84 | 72.24 | 0 | 0 | 0 | 25.92 | 43.35 | 0.15 | 2586.98 |

Appendix

apA

**Table B.** Group of reach variables measured *in situ.* For the substrate type we calculate a visual estimation in every sampled reach.

| Variable Category | Variable | Unit | Definition |
|---|---|---|---|
| Geomorphological | | | |
| | Distance to the lake | Km | Measured with the google earth tool from the highest part of the sampled section to the mouth |
| | Length of the main flow | Km | length of the main channel measured from the source to the mouth of the stream |
| | Length of the Valley | Km | length of the valley from the stream header to the mouth |
| | Wetted Width | mts | transect width with water |
| | Active -channel Depth | cm | water depth of the active channel |
| | Active- channel Width | mts | The width of the active channel measured perpendicular to streamflow. The active-channel, a short-term geomorphic feature formed by prevailing stream discharges, is narrower than the bankfull channel and is defined by a break in bank slope that also typically is the edge of permanent vegetation |
| | Bankfull Depth | cm | the difference in elevation between the bankfull stage and the deepest part of the cross section |
| | Bankfull Width | mts | The width of the bankfull channel measured at a section perpendicular to streamflow at bankfull discharge |
| | Width/Depth Ratio (W/D) | | is the bankfull surface width divided by the bankfull mean depth. The width/depth ratio describes the channel shape (large number= wide and shallow; small number= narrow |

| | | | |
|---|---|---|---|
| | | | and deep) |
| | Entrenchment Ratio (ER) | | Entrenchment is the vertical containment of a river and is quantitatively defined as the width of the flood-prone area divided by the bankfull surface width |
| | Slope | % | Rise/run x 100 Where the rise/run value is: (highest elevation-lowest elevation)/distance between elevation points |
| | Sinuosity | | ratio of stream channel length to basin length |
| Substrate type | Pool proportion | % | |
| | Silt | % | |
| | Sand | % | |
| | Gravel (2-64mm) | % | |
| | Cobble (64-256MM) | % | |
| | Boulder (>256mm) | % | |
| | Bedrock | % | |
| Other habitat variables | Stream order | | *sensu* Bain and Stevenson (1999) |
| | Riparian Forest | | Considering the dominant species |
| | Woody debris | Number of pieces | number of wood pieces (>1m lenght) within the channel |
| | Land Use | | Type of land use registered near the reach |

### Appendix

apA

| | | | |
|---|---|---|---|
| | Benthos | | Benthos biomass collected with a surber sampler (0.09 m2; 250 µm pore size) |
| Hidrological | | | |
| | Flow velocity | m/s | rate of water movement |
| | Discharge | $m^2/s$ | volume of water passing a point per unit time: calculated with a program |
| Water quality | | | |
| | Turbidity | NTU | Field Nephelometer |
| | Temperature | °C | |
| | Conductivity | µS/cm | |
|  | pH | pH | All these variables were calculated with a field multiparameter probe |
| | Total Dissolved Solids | mg/l | |
| | Salinity | ppt | |

apA

**Table C.** CPUE (number of fish / 100m$^2$) in the 35 watersheds of the Limay river basins.

| Watershed | Latitude | Longitude | CPUEN | | | |
|---|---|---|---|---|---|---|
| | | | *O. mykiis* | *S. trutta* | *S. fontinalis* | Total |
| 3.Pireco | -40.7283365 | -71.883447 | 0.76 | 0.69 | - | 1.45 |
| 2.Acantuco | -40.6870432 | -71.8259462 | 1.63 | 0.4 | 0.04 | 2.07 |
| 4.Machete | -40.837265 | -71.8331876 | 4.01 | 4.48 | 0.24 | 8.73 |
| 5.Gallardo | -40.8700759 | -71.8211752 | 1.75 | 5.5 | - | 7.25 |
| 10.Blest | -41.0243555 | -71.8452124 | 9.5 | 1.6 | 0.32 | 11.43 |
| 11.Patiruco | -41.0651909 | -71.7490861 | 12.13 | - | 2.55 | 14.69 |
| 12.Frey | -41.1711721 | -71.7299522 | 1.76 | 1.63 | 0.26 | 3.78 |
| 9.Millaqueo | -40.9742568 | -71.6598227 | 11.01 | - | 0.35 | 11.36 |
| 6.Coluco | -40.9123437 | -71.6687729 | 3.93 | 2.02 | 0.12 | 6.07 |
| 1.Neuquenco | -40.5767528 | -71.6595039 | 5.16 | 1.54 | 0.09 | 6.78 |
| 14.Lluvuco | -41.145673 | -71.611091 | 1.34 | 0.11 | - | 1.45 |
| 25.Casa de Piedra | -41.1604288 | -71.5157383 | 11.27 | - | 0.17 | 11.44 |
| 15.Bonito | -40.7356713 | -71.5787519 | 14.88 | 0.56 | 0.28 | 15.73 |
| 16.Estacada | -40.7829627 | -71.5257495 | 1.88 | - | - | 1.88 |
| 17.Ragintuco | -40.8125503 | -71.4787445 | 1.1 | - | - | 1.1 |
| 18.Huemul | -40.8569227 | -71.4418746 | 7 | - | - | 7 |
| 19.Pedregoso | -40.9034006 | -71.3690179 | 1.45 | - | - | 1.45 |
| 20.Quintriquenco | -40.9247984 | -71.3205718 | 8.19 | - | - | 8.19 |
| 24.Castilla | -41.022637 | -71.3417586 | 10.01 | 36.48 | 7.72 | 54.21 |
| 21.Manzano-Jones | -40.9812028 | -71.2790299 | 17 | 9.46 | - | 26.46 |
| 23.Chacabuco | -40.9954146 | -71.2287048 | 3.91 | 13.78 | - | 17.69 |
| 31.Tristeza | -41.2894088 | -71.3208816 | 20.56 | 0.32 | 0.32 | 21.19 |
| 26.Cascada | -41.1560917 | -71.4530239 | 39.7 | - | - | 39.7 |
| 27.Gutiérrez | -41.2066666 | -71.4325938 | 1.05 | 7.64 | - | 8.69 |
| 30.Torrontegui | -41.2788083 | -71.4389803 | 22.61 | 1.19 | 0.59 | 24.39 |
| 28.Ñireco | -41.2078291 | -71.321543 | 18.8 | 0.09 | - | 18.89 |
| 32.Challhuaco | -41.2365767 | -71.3090935 | 42.43 | - | 2.86 | 45.29 |
| 35. de la Quebrada | -41.3616476 | -71.271492 | 50 | 2.14 | 0.43 | 52.56 |
| 29.del Medio | -41.1808270 | -71.2128754 | 20.56 | 5.76 | - | 26.32 |
| 34.Las Minas | -41.2928298 | -71.1703178 | 18.86 | 0.64 | - | 19.5 |

| | | | | | | |
|---|---|---|---|---|---|---|
| 33.Ñirihuau | -41.2252532 | 71.1862508 | 8.92 | 0.54 | - | 9.47 |
| 7.Bravo | -40.9686895 | -71.8035434 | - | - | - | - |
| 8.Blanco | -40.9862577 | -71.7268875 | - | - | - | - |
| 22.Ñewbery | -40.9800624 | -71.1866559 | - | - | - | - |
| 13.Uhueco | -41.1670992 | -71.6556875 | - | - | - | - |
